# A Simple Optimization Workflow to Enable Precise and Accurate Imputation of Missing Values in Proteomic Datasets

**DOI:** 10.1101/2020.07.24.220467

**Authors:** Kruttika Dabke, Simion Kreimer, Michelle R. Jones, Sarah J. Parker

## Abstract

Missing values in proteomic data sets have real consequences on downstream data analysis and reproducibility. Although several imputation methods exist to handle missing values, there is no single imputation method that is best suited for a diverse range of data sets and no clear strategy exists for evaluating imputation methods for large-scale DIA-MS data sets, especially at different levels of protein quantification. To navigate through the different imputation strategies available in the literature, we have established a workflow to assess imputation methods on large-scale label-free DIA-MS data sets. We used two distinct DIA-MS data sets with real missing values to evaluate eight different imputation methods with multiple parameters at different levels of protein quantification; dilution series data set and an independent data set with actual experimental samples. We found that imputation methods based on local structures within the data, like local least squares (LLS) and random forest (RF), worked well in our dilution series data set whereas, imputation methods based on global structures within the data, like BPCA performed well in our independent data set. We also found that imputation at the most basic level of protein quantification – fragment level-improved accuracy and number of proteins quantified. Overall, this study indicates that the most suitable imputation method depends on the overall structure and correlations of proteins within the data set and can be identified with the workflow presented here.

## Introduction

Mass spectrometry is a powerful tool to rapidly identify and quantify thousands of peptides and proteins in a wide variety of experimental designs and disease specimens^1–3^. Studies by Clinical Proteomic Tumor Analysis Consortium (CPTAC)^4–6^, Human Tumor Atlas Network (HTAN)^7^, Cancer Cell Line Encyclopedia (CCLE)^8^ among others, have used class prediction and class discovery methodologies, to understand how protein expression changes correlate to the presence of disease. To grasp the underlying biological reality based on protein expression profiles, it is essential to eliminate artifacts like noise and to maximize data usability by managing missing values generated as a consequence of various steps and data filtering steps along the analytical pipeline. Although powerful, multivariate analysis tools like hierarchical clustering or principal component analysis often have difficulty treating missing values. Different ways of dealing with missing values often lead to variations in outcome^9^. Thus, understanding the nature, extent, and ways to handle missing values is a crucial preprocessing step in data analysis.

In mass spectrometry studies, missing values are defined as intensities or values not recorded for a peptide. In LC-MS/MS approaches, the range of missing values can be from 10% to 50% overall, while a very high proportion of peptides/proteins can have at least one missing value (70% - 90%)^10^. Missing values are generated if: 1) the peptide intensity was below the detection limit of the instrument, 2) data analysis algorithm failed to detect the peptide, or 3) the peptide was not present in the sample. There could be other reasons for missing values like mis-cleavage, ionization competition, ion suppression or peptide misidentification^11^. Due to the nature of protein intensity quantification, missing values (or values recorded as NAs) introduced at the peptide level are aggregated to protein intensities and can cause issues with hypothesis testing, reproducibility, and consistency of proteomic data analysis. Advances in the proteomics field in the past few years have led to a significant improvement in reproducible protein quantification, especially with Data Independent Acquisition mass spectrometry (DIA-MS)^12–14^. DIA-MS allows for improved sensitivity and accuracy in protein quantification which translates to fewer missing values. However, the missing values are not eliminated with this state-of-the-art technology. DIA-MS data analysis, including scoring of peptide identification, retention time alignment, and calculating false discovery rates across a large number of samples, can be a significant contributor to introducing missing values at various steps of the analysis pipeline^15^.

Whatever the reason for missing values, they can statistically be divided into three categories^11,16^:

1. MCAR - Missing completely at random can best be described in proteomics as an accumulation of multiple minor errors leading to missing protein intensity where the missingness of a peptide can neither be explained by the nature of the peptide or its abundance.
2. MAR - Missing at random means there is a relationship between the propensity of missing values and observed data of an individual’s variable. It is a more general definition of MCAR values. Both MCAR/MAR are considered the same in proteomics data sets, and both are randomly distributed.
3. MNAR-Missing not at random in proteomics data sets are values that are missing because they are below the limit of detection of the instrument and are usually called the “non-ignorable” missing values.

Merely ignoring missing values of any nature could lead to a decrease in size or completeness of data, limiting the researcher’s ability to interpret peptide/protein concentrations. Imputation has become an important tool in handling missing values. A variety of imputation algorithms have been developed in the past but they mainly fall under three categories; imputation based on 1) single-digit replacement (zero, minDet, minProb imputation) 2) local structures in datasets (knn, lls) 3) global structures in datasets (BPCA)^17^. Although developed for microarray-based analysis, these methods have been explored in proteomic data sets. A study in mass spectrometry-based metabolomics data found that random forest-based method of imputation performed best in case of MCAR/MAR type missing values and quantile regression imputation of left-censored data (QRILC) imputation method worked best for MNAR type missing data^18^. Another study that compared five different imputation methods in label-free proteomics data found that local least square (lls) regression imputation paired with filtration strategies worked best to identify differentially expressed proteins accurately^19^. Taken together, these findings suggest that optimization analyses to determine the ideal imputation strategy for a given experiment or data structure is advisable.

Questions regarding how to structure an optimization experiment to best evaluate imputation performance, what metrics to monitor, and at what data level to perform imputation (e.g., fragments, peptides, or proteins) remain unclear. A group recently developed a software called NAguideR to impute missing values for DIA-MS data sets. They assessed 23 imputation methods with six criteria to evaluate their performance^15^. Studies described above evaluated imputation methods using simulated missing values.

They took a complete data set and randomly generated different percentages of MCAR/MAR and MNAR missing values. This does not entirely reflect the exact pattern of missing values present in a real data set and pattern of real missing values is hard to define or simulate accurately. Another study evaluated nine imputation methods using proteomic data sets with real missing values^17^, but they did not explore imputation of missing values at the fragment level. Fragment and/or protein level imputation could lead to increased accuracy and numbers of proteins quantified, as missing peptides from one sample to the next can strongly impact estimates of protein level intensities. Therefore, a study looking into imputation of real missing values at different levels of protein quantification for DIA-MS data sets is currently missing.

In this study, we present a workflow to test and evaluate imputation strategies on DIA-MS proteomic data, utilizing an initial loading dilution series as an optimization dataset along with the inclusion of replicate acquisitions of selected individual biological samples in order to enable evaluation of different imputation approaches and selection of a preferred approach for a given data structure and experiment. We evaluated eight commonly used imputation methods on a dilution series data set with real missing values. We assessed the performance of imputation methods using multiple parameters at different levels of protein quantification, i.e., fragment, peptide, and protein. We then applied the top imputation strategies from the dilution series to an independent dataset comparing actual experimental samples, as well as technical replicates of pooled lysate. We present our findings and this workflow as a data acquisition strategy to enable analytical optimization and data QC for large-scale, label-free DIA-MS studies.

## Results

### Evaluation of the dilution series data set used for imputation analysis

We generated a dilution series to be used as an optimization data set for evaluating imputation methods. It was generated by pooling protein lysates from all individual patient samples and loading three different concentrations of 2,4 and 8 ugs (referred to as low, medium, and high load respectively) for LC MS/MS in triplicate (*Figure 1.A*). With this approach, we quantified 2918 proteins in at least one of the nine MS data files. As expected, we observed an inverse correlation between the percentage of missing values and total peptide load used as input for MS (bar plot annotation of the heatmap, *Figure 1.B*). Unsupervised hierarchical clustering of the replicates (*Figure 1.B*) showed replicates clustering by the load. Pearson’s correlation coefficient calculated for each replicate pairwise correlation showed a high correlation of replicates within a load compared to replicates across loads (*Figure 1.C*). Since the protein lysates were loaded on to MS in increasing concentration, we saw the expected stepwise increase in median protein intensity across different loads with a median log-2 fold protein intensity of 11.92, 10.99, and 9.78 for high, medium and low load respectively (*Figure 1.D*). As expected from serial injections of the same sample digest, we observed generally low coefficient of variance across the three MS replicates within each load (median coefficient of variance = 10.68, 9.08, 8.89 for high, medium, and low load, *Figure 1.E*). The dilution series data set showed expected trends in protein intensities and replicate correlations which made it appropriate for use in further evaluation of imputation methods.

**Figure 1:**
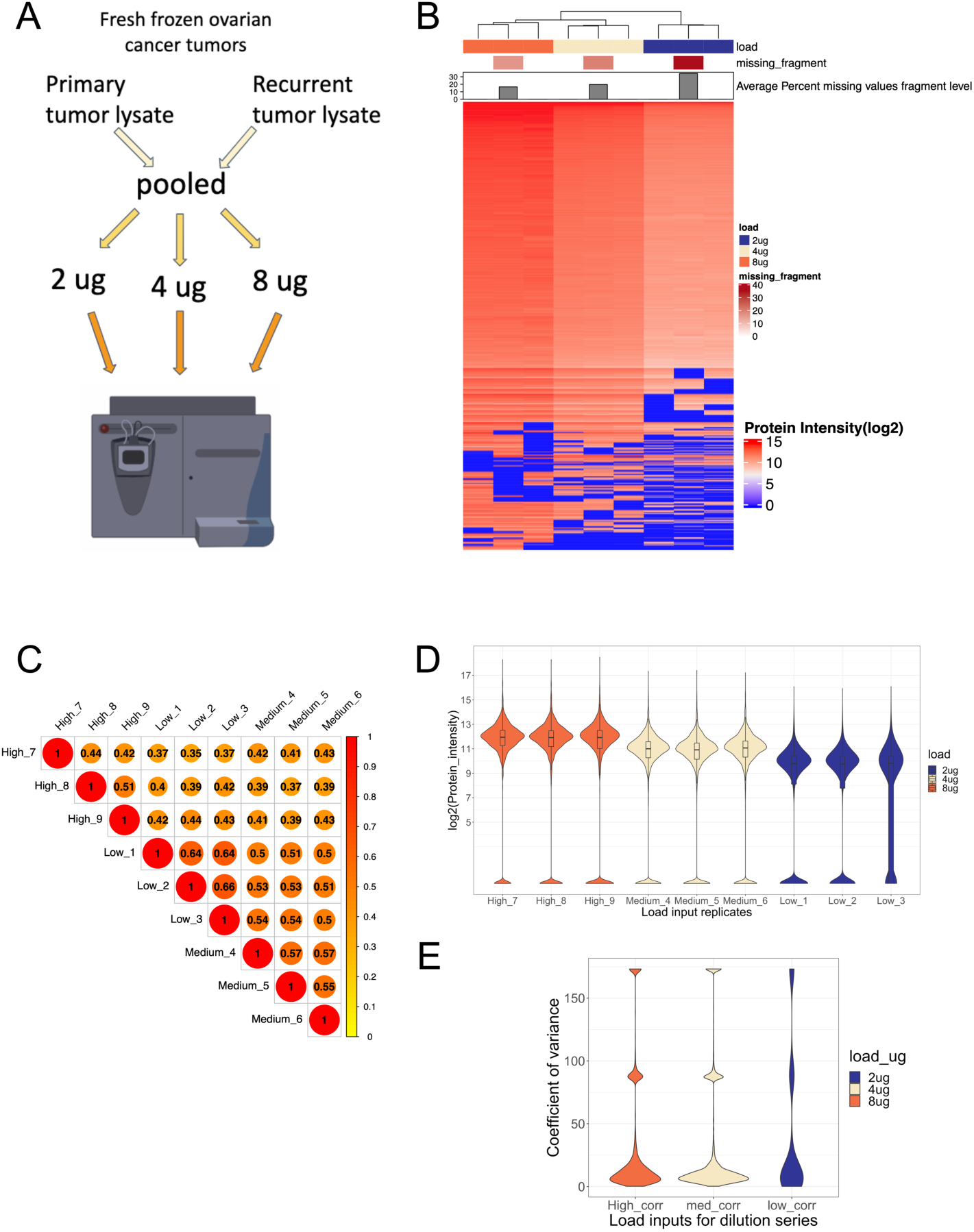
Comprehensive evaluation of the dilution series data set. **A)** Schematic of the dilution series experiment. **B)** Unsupervised hierarchical clustering of protein intensities (rows) shows clustering of samples (columns) by replicates. **C)** Pearson’s correlation coefficient of replicates by protein intensities shows a high correlation among all replicates within loads (all correlations are significant; p-value < 0.001) **D)** Distribution of protein intensities of replicates show a step wise decrease in median protein intensity corresponding to decreasing load **E)** Coefficient of variance (CV) calculated by protein intensities of the 3 replicates within each load.

### Analysis of imputation methods

We tested eight imputation methods to handle missing data in the optimization data set described above. These methods were chosen because they covered the three categories of missing value imputation methods:

1. Single-digit replacement (zero, minDet, minProb)
2. Local structures in datasets (knn, lls, rf)
3. Global structures in datasets (svd, bpca)

We performed imputation at different levels of protein quantification i.e., fragment, peptide, and protein levels. A total of 182,410 (23.52%), 22,130 (19.83%), 4,187 (15.94%) missing intensities across all nine replicates were subsequently imputed at the fragment, peptide and protein levels respectively. Due to the nature of the dilution series data set, for which a specific quantitative difference between data groups has been introduced, we were able to use an ideal/expected log fold change of protein intensities between different loads to measure how well the imputation methods performed. *Figure 2.A* shows the distribution of pairwise mean log fold change between different loads for the eight imputation methods compared to the non-imputed data set. Two horizontal lines show the expected (e.g., ideal) log2 fold change for the pairwise fold change calculations. Of the eight methods analyzed, lls and random forest imputation methods had the smallest spread around the expected mean values (e.g., Log2FC of 1 or 2) for pairwise comparisons for each protein between any two dilution values, indicating that following imputation the precision of quantification by these methods was best for these data. *Figure 2.B* shows that random forest and lls based imputation methods have the highest Further, Pearson’s correlation coefficients with the expected/ideal log fold change (0.94 and 0.9, respectively). We also calculated the ratio of RMSE/S.D. within the fold-change calculations before and after imputation where knn, lls, and random forest imputation (RF) methods showed the least error compared to expected/ideal fold change (0.83,0.54 and 0.43 respectively, *Figure 2.C*). To assess whether the level (e.g., fragment, peptide or protein) at which imputation is performed affects quantitative precision, we chose the top three best performing imputation methods shown in *Figure 2*, i.e. knn, lls and random forest, and also performed imputation at the peptide and protein levels (*Supplementary Figure 1*). We calculated Pearson’s correlation coefficient for the three imputation methods at peptide (*S.Figure. 1.C*) and protein levels (*S.Figure 1.D*) along with error estimations for the two levels of imputation compared to expected/ideal fold changes (peptide level – *S.Figure 1.E* and protein level – *S.Figure 1.F*). Fragment level imputation subtly but consistently outperformed the peptide and protein level imputation. Fragment level imputation also resulted in slightly larger protein ID count, 2918 versus 3026 protein IDs quantified after fragment and peptide/protein level imputation respectively.

**Figure 2:**
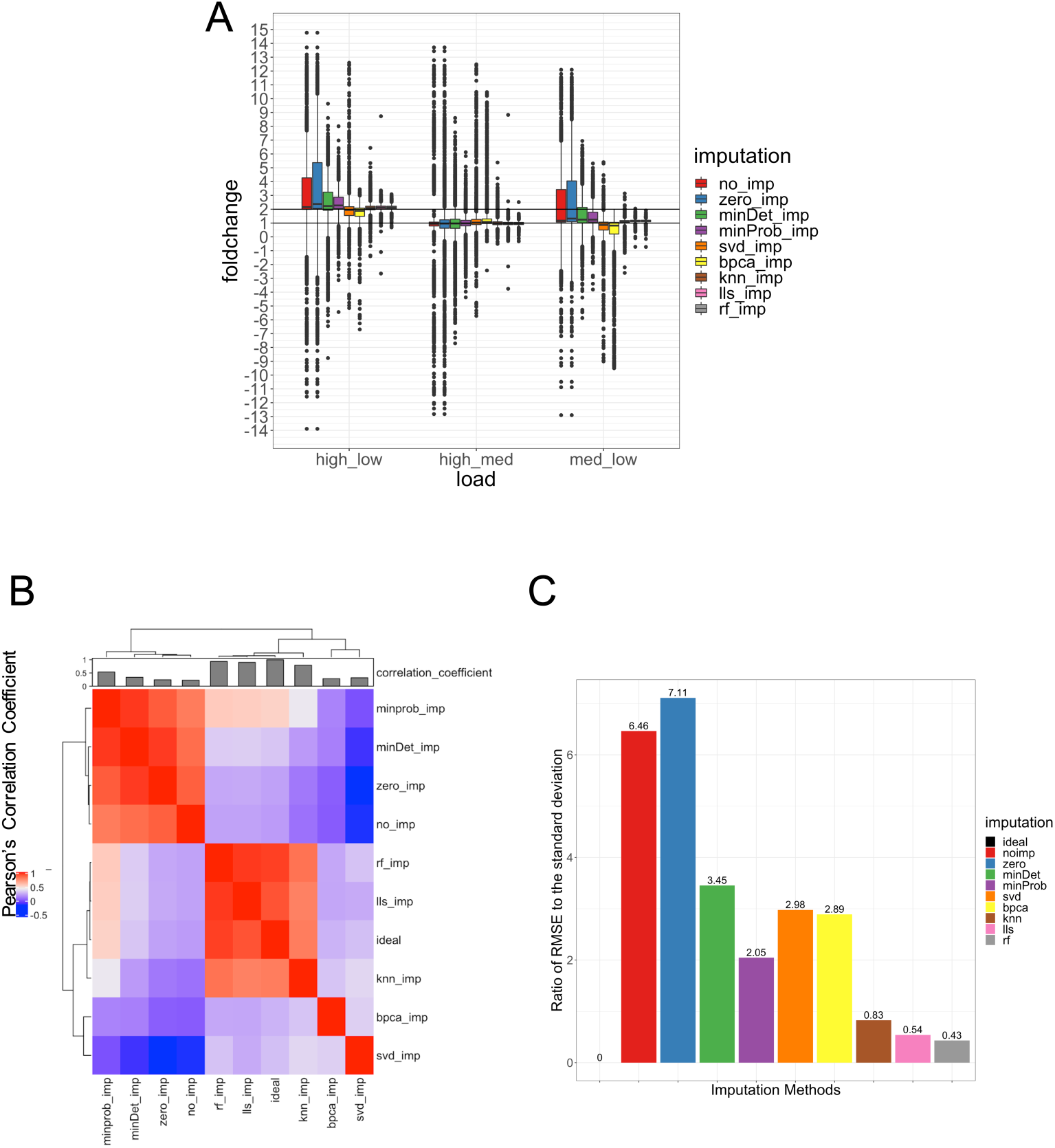
Analysis of imputation methods; imputation at the fragment level. **A)** Pairwise fold changes calculated across three loads for different imputation methods shows values from local least squares (lls) and random forest imputation closest to expected fold change with the least number of outliers **B)** Hierarchical clustering of Pearson’s correlation coefficient shows clustering of fold change after lls and random forest imputation with ideal/expected fold change **C)** Ratio of root mean square error (RMSE) to standard deviation (SD) between fold changes of different imputation methods and ideal/expected fold change shows the least error for lls and random forest imputation methods.

In addition to the analysis of quantitative performance, we further evaluated the dilution series data set using the NAguideR tool^15^, which provides a classic and proteomic criterion for evaluation of imputation methods for proteomic data sets. The classic criteria calculate normalized root mean square error (NRMSE), NRMSE based sum of ranks (SOR), the average correlation coefficient between the original and imputed values, and Procrustes statistical shape analysis (PSS). The proteomic criteria calculate the average correlation coefficient within each protein’s corresponding peptides, average correlation coefficient within every protein complex based on the CORUM database, and average correlation coefficient within each cluster of protein-protein interaction network based on hu.MAP database. The tool takes peptide level intensities with corresponding protein ID labels as input and requires a maximum missingness filter. We filtered the dilution series data set such that each peptide did not have more than 80% missing values, which resulted in a total of 9,916 (80%) peptides being retained for analysis. Bpca, lls, and rf based imputation methods performed the best as per the classic criteria (*Figure 3.A*) while bpca, knn, and lls based imputation methods performed the best as per the proteomic criteria (*Figure 3.B*). Our analysis showed that the random forest-based imputation method had the least variation in pairwise log fold changes and the lowest error estimations compared to ideal fold changes when imputation was performed at the fragment level. Peptide level imputation resulted in bpca, lls, and knn imputation methods performing the best based on classic and proteomic criteria calculated using the NAguideR tool.

**Figure 3:**
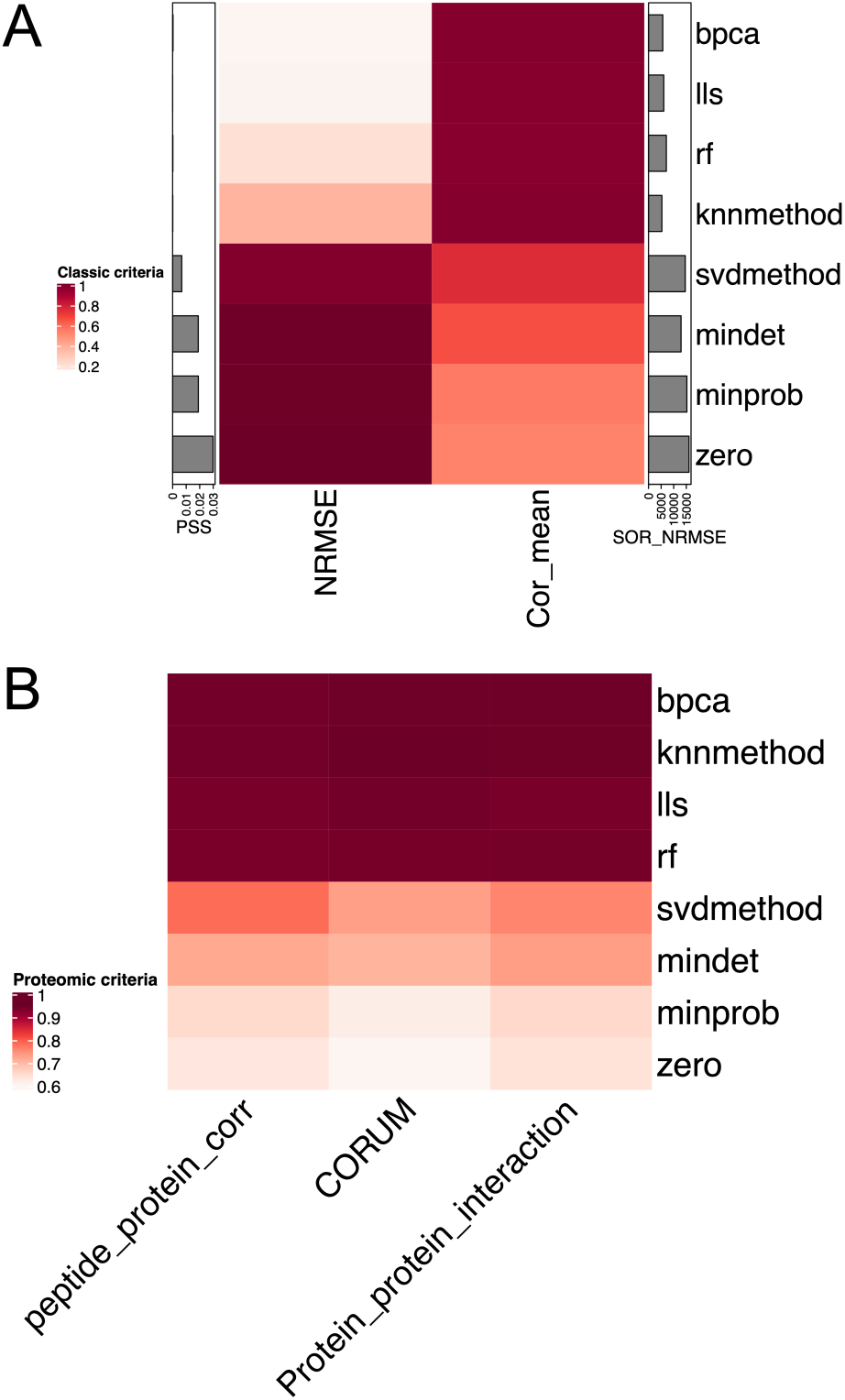
Evaluation of imputation methods using the NAguideR tool after imputation at peptide level. **A)** Classic criteria for evaluation of imputation methods at peptide level. (PSS = Procrustes statistical shape analysis, NRMSE = Normalized root mean square error, Cor_mean = Average correlation of the entire data set, SOR_NRMSE = Sum of ranks of NRMSE). **B)** Proteomic criteria for evaluation of imputation methods at peptide level. (peptide_protein_corr = Average correlation within each protein’s corresponding peptides, CORUM = average correlation coefficient within every protein complex based on the CORUM database, Protein_protein_interaction = average correlation coefficient within each cluster of protein-protein interaction network based on hu.MAP database)

### Summary of the dilution series data set after imputation of missing values with random forest-based imputation

Our evaluation signified that the imputation of missing values at the fragment level with random forest-based imputation worked best compared to eight other imputation methods for the dilution series data set. We assessed the effects of fragment level random forest-based imputation on the data structure, in the context of protein intensities. Random forest imputation preserved clustering (*Figure 4.A*) and improved correlations of replicates (*Figure 4.B*) within each load (scale of Pearson’s correlation coefficient changed from 0.3-1 to 0.94-1 for non-imputed vs. imputed data sets). Random forest imputation of missing values conserved the stepwise increase in median protein intensity, which was inherent to the dilution series data set (*Figure 4.C*). The coefficient of variance for all replicates within loads decreased after imputing missing values using random forest-based imputation (*Figure 4.D*). Thus, random forest-based imputation of missing values at the fragment level did not change the dataset’s inherent properties but further improved correlation of the replicates for each load.

**Figure 4:**
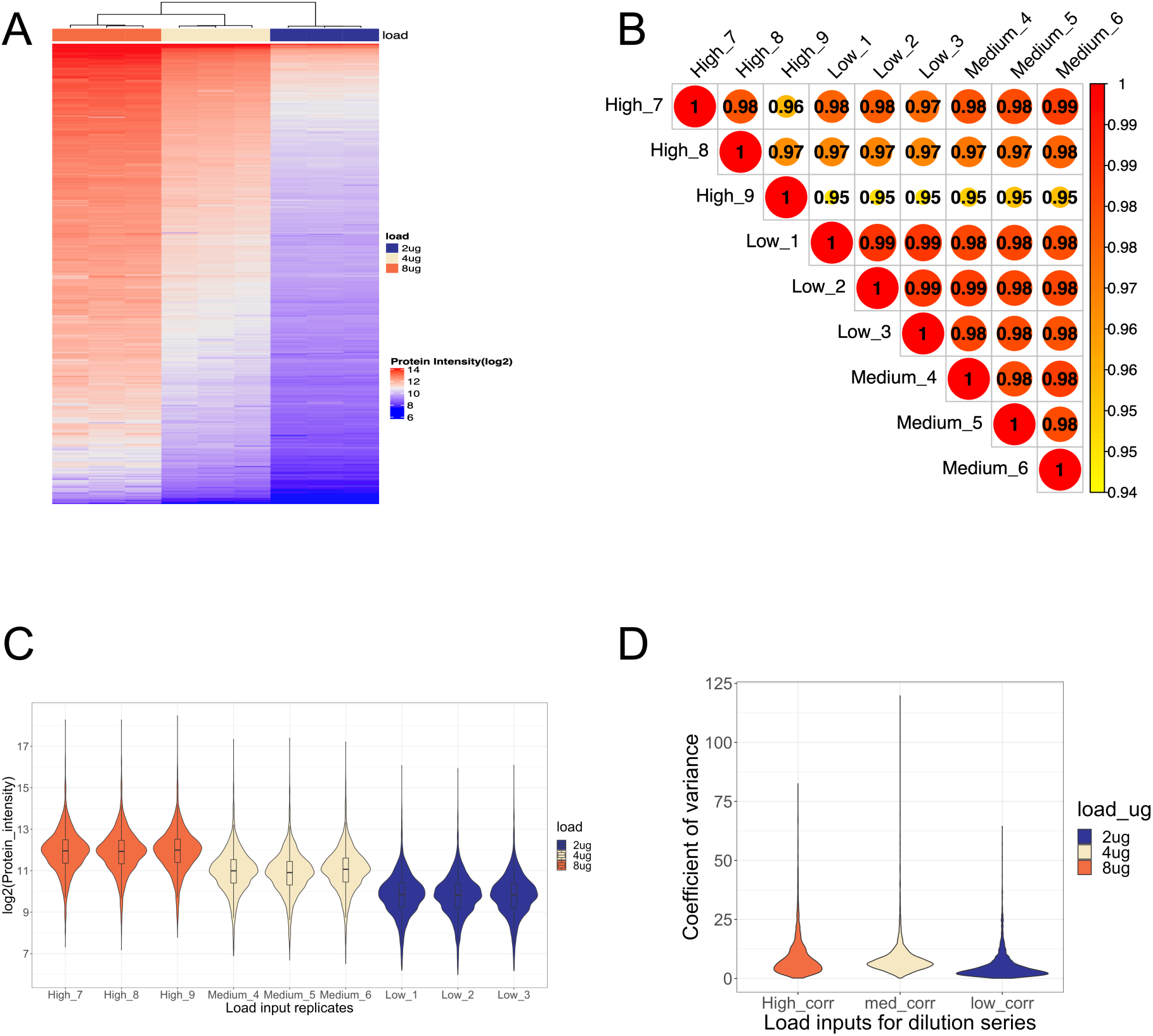
Evaluation of the dilution series data set after imputation of missing values using random forest. All figures are plotted after nonparametric missing value imputation using random forest **A)** Unsupervised hierarchical clustering of protein intensities (rows) shows clustering of samples (columns) by replicates **B)** Pearson’s correlation coefficient of replicates by protein intensities shows a high correlation among all replicates across different loads (all values are significant; p-value < 0.001) **C)** Distribution of protein intensities of replicates show a step wise decrease in median protein intensity corresponding to decreasing load **D)** Coefficient of variance (CV) calculated by protein intensities of the 3 replicates within each load.

### Analysis of imputation methods on an independent data set

To further evaluate the findings discussed above, we tested the eight imputation methods on an independent data set. The independent data set consisted of protein lysates from six individual ovarian cancer tumors analyzed by MS in duplicates or triplicates on a different MS set up (e.g., Orbitrap LUMOS versus TripleTOF 6600 for dilution). We quantified 5410 proteins across fourteen samples, including missing values. A total of 1,082,702 (49.14%), 138,271 (43.84%), and 25,278 (33.33%) missing intensities were imputed at the fragment, peptide and protein levels respectively across all replicates. Pearson’s correlation coefficient calculated based on protein intensities showed a correlation among replicates of the same sample between 0.7-0.9 depending on sample (*S.Figure 2.A*). To test the performance of eight imputation methods, we used the NAguideR tool, which provides a classic and proteomic criterion for evaluation. The tool takes peptide and protein level intensities as input. For this study, the independent data set was filtered such that each peptide across all samples did not have more than 80% missing values, which resulted in 14,418 (64%) of the data set being retained for analysis. The top-ranking imputation methods were lls and rf as per the classic criteria (*Figure 5.A*) while lls, bpca, and rf based imputation methods performed the best as per the proteomic criteria (*Figure 5.B*). We calculated the average Pearson’s correlation within each sample’s corresponding replicates, before and after imputation, and observed that some imputation approaches improved while others were detrimental to the agreement between technical replicates of the same sample. *Figure 5.C* shows that random forest-based imputation had the highest average correlation per sample compared to six other methods (*S.Figure 2. B-G*). Random forest-based imputation also decreased the coefficient of variance of replicates within samples compared to the coefficient of variance without imputation (*Figure 5.D, E*). Overall, imputing missing values improved the correlation of replicates within samples and decreased their coefficient of variance.

**Figure 5:**
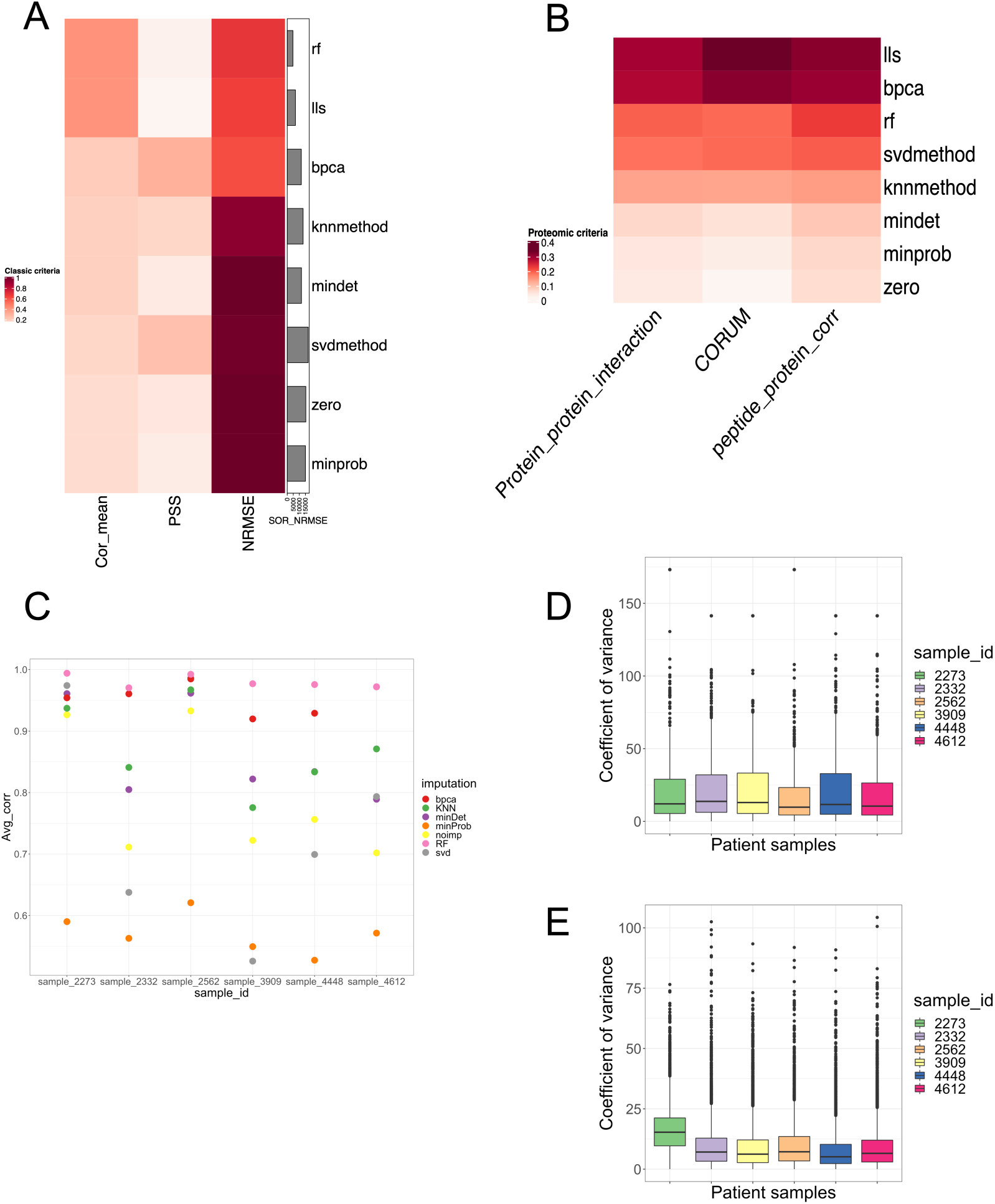
Analysis of imputation methods on an independent data set. **A, B)** Evaluation of imputation methods after peptide level imputation using classic (A) and proteomic criteria (B) from the NAguideR tool. **C)** Average Pearson’s correlation coefficient within samples of the independent data set shows improved correlation after missing value imputation with random forest and bpca. **D, E)** Coefficient of variance within samples before (D) and after (E) imputation with random forest-based imputation method.

## Discussion

Missingness due to a multitude of potential causes is a persistent issue plaguing discovery proteomics experiments. Missingness at the level of fragments and in particular peptides of proteins within a dataset has the potential to be particularly detrimental due to its impact on the estimation of a given protein abundance level between samples of a dataset. While stringent approaches such as only including peptides seen in all samples for protein abundance estimates are an option, this will lead to loss of potentially valuable quantitative information for each protein as well as reduced number of quantifiable proteins. In this study, we have explored the effect of a handful of different approaches for the imputation of missing values on the quantitative and overall fidelity of biological proteomic data. Importantly, we performed and tested the effect of imputation at the most raw level of quantification - the peptide fragments - of a DIA dataset. We found that imputation methods based on the local structure of the data (e.g., LLS and RF) tended to outperform all other methods, and that while imputation at fragment level relative to peptide or protein level imputation only marginally improved overall quantitative precision in our dilution series dataset, it resulted in a larger number of quantifiable proteins within the experiment overall.

Interestingly, while the local structure-based imputation clearly outperformed other methods on the experimentally produced gradient dataset, the global structures imputation approach (bpca) had demonstrably better performance on the individual dataset relative to its performance on the gradient dataset. These differences in performance may be explained by the inherent properties of the proteomic data sets.

Strong local correlations were present in the dilution series data set due to experimental design. The independent data set on the other hand, had much higher protein complexities with unknown drivers of protein expression, making local structures hard to define and thus methods like BPCA which look at global covariance structures among samples/proteins performed well. Even within the local structures, RF and LLS outperformed KNN method of imputation. One reason could be that KNN simply looks at k nearest neighbors (where k = 10-20) to impute missing values which in the case of gene expression studies are good representatives of missing values since they are co-expressed frequently and are quantified in large numbers (> 10,000). In contrast, proteomic data sets do not generally have significant co-expression patterns due to relatively smaller numbers of proteins quantified per sample. RF and lls also assume strong local correlations within expression matrix but impute missing values with a more advanced iterative approach, which is probably why they performed better than other local structure-based imputation methods. Single value methods are simple and quick to implement in large scale data sets, but they generally work well only for MNAR type missing values and tend to severely bias the data which is why these methods tend to perform poorly.

Overall, these results demonstrate that inclusion of 1) a dilution series of sample pool and 2) technical replicates within a biological dataset can both be leveraged to inform the ideal approach for data imputation in a given experiment and biological data structure. While technical replicates of every sample may be inefficient, especially for very large datasets, a random subset of replicates can suffice as a read out. Alternatively, the use of regularly spaced digestion replicates of a samples pool could be used to inform best imputation method for a given experiment. In this experiment, the RF imputation method tended to perform best across most, but not all, indicators with the LLS and BPCA methods also performing well. We do not believe that these methods will be the top choice for all datasets, however, and thus present our findings as an example of simple approaches that can be added to any quantitative discovery proteomics workflow with minimal extra sample requirements or MS time demands. To summarize, in this paper we have outlined a workflow and system of metrics, including the recently published NAguideR indicator suite, to evaluate and select an imputation approach that is appropriate for a given biological proteomic dataset.

## Material and Methods

### Preparation of fresh frozen tissue

The Women’s Cancer Program Biorepository at Cedars-Sinai Medical Center has collected extensive clinical data and specimens from more than 16,000 patients over 25 years. For this study, we used a set of 12 HGSOC patients diagnosed with Stage III/IV disease for whom paired primary (chemo-naïve) and recurrent (chemoresistant) tumors were available.

Fresh frozen tumors were embedded in optimal cutting temperature compound, bisected, and mounted before slides were made for hematoxylin and eosin staining. All slides were reviewed by a single pathologist to identify regions of epithelial carcinoma. A 3mm core of approximately 50mg was then extracted from a region of pure epithelial carcinoma from the intact frozen tumor, with care not to include any of the surrounding stroma. These ‘pure tumor’ tissue cores were used for subsequent proteomic analyses. Frozen tumors were thawed on ice, 200 – 400 ul lysis buffer (6M Urea + 0.1% Rapigest) was added, and tissue was homogenized with a polytron. Proteins were extracted by high-pressure barocycling on a Pressure BioSciences instrument, model 2320EXT (PBI is Easton, MA) at room temperature, ramping pressure to 45 psi, holding for 50 s, returning to atmospheric pressure for 10 s, and repeating for 60 cycles. Protein concentration was assayed using the Pierce BCA assay. Two separate datasets were then prepared from the homogenates, a pool of all patient samples as well as a small pilot group of individual patient samples.

For the dilution series data set, we pooled 20ug of protein lysates from each sample. The pooled protein lysate was diluted to <2M urea using 50mM ammonium bicarbonate with 50mM DTT added as a reducing agent for 30 min at 370C. Protein lysate was alkylated with 50mM iodoacetamide for 30 min at room temperature in the dark to alkylate reduced thiol groups. 1ug of Trypsin/Lys-C Mix: 50ug of protein lysate was used to digest reduced and alkylated protein lysate at a pH of ∼7-8. The lysate was digested with high-pressure barocycling at 370C with the following settings: 45kPSI, 60 cycles, 50s on, 10s off, and maintained overnight at 370C. After digestion, peptides were desalted with NEST desalting tips and eluted off the column using 50% Acetonitrile: 50% of 0.1% formic acid in water. A BCA analysis of peptide concentration was performed on the eluted, desalted peptides. The peptides were then dried and stored at -20C. The dried sample was reconstituted with 0.1% formic acid in water diluted to 2,4, and 8 ug (referred to as low, medium, and high load respectively) before acquisition on MS.

For the individual patient data set was acquired from 6 l patient ovarian cancer tumors (3 of each group) digested into peptides and desalted as described above. Importantly, the individual patient samples were processed and acquired at different time points, making it independent from the previous analysis. At least 2 technical replicates of each individual sample was acquired in order to assess the impact of different imputation methods within a given sample.

### Data-Independent Mass Spectrometry

#### Data acquisition; dilution series data set

2,4 and 8 ugs of peptide were injected in triplicate on to spiked with exogenous retention time peptide standards (iRT, Biognosys) and loaded onto an Eksigent 415 HPLC system via an Ekspert nanoLC 400 autosampler. Peptides were first loaded onto a trap column (10×0.3 mm, C18CL, 5 µm, 120Å, Sciex) for 3 minutes at 10 µL/min of solvent A (0.1% FA in water) the separated on an analytical column (ChromXP C18CL, 150×0.3 mm, 3µm, 120Å, Sciex) at a flow rate of 5 µL/min using a linear AB gradient of 3-35% solvent B (0.1% FA in ACN) for 60 minutes, 35-85% B for 2 minutes, holding at 85% B for 5 minutes, then re-equilibrating at 3% B for 7 minutes. Mass spectra were collected in data independent acquisition mode, with an initial MS1 scan of 250 ms ranging from 400-1250 m/z followed by the acquisition of 100 variable width MS2 scans of 30 ms, collecting data on ions between 100-1800 m/z. Variable window set up has been published previously^20^. Total cycle time was 3.3 seconds. Source gas 1 was set to 15, gas 2 was set to 20, curtain gas set to 25, source temperature set to 100 °C, and source voltage set to 5500 V.

#### Data acquisition; independent data set

4 uL of digested sample were injected directly unto a 200 cm micro pillar array column (uPAC, Pharmafluidics) and separated over 120 minutes reversed phase gradient at 1200 nL/min and 60 C. The gradient of aqueous 0.1% formic acid (A) and 0.1% formic acid in acetonitrile (B) was implemented as follows: 2% B from 0 to 5 min, ramp to 4% B at 5.2 minutes, linear ramp to 28% B at 95 minutes, and ramp to 46% B at 120 minutes. After each analytical run, the column was flushed at 1200 nL/min and 60 C by injection of 50% Methanol at 95% B for 25 minutes followed by a 10 minutes ramp down to 2% B and a 5 minute equilibration to 2% B.

The eluting peptides were electro sprayed through a 30 um bore stainless steel emitter (EvoSep) and analyzed on an Orbitrap Lumos using data independent acquisition (DIA) spanning the 400-1000 m/z range. Each DIA scan isolated a 15 m/z window with no overlap between windows, accumulated the ion current for a maximum of 54 seconds to a maximum AGC of 5E5, activated the selected ions by HCD set at 30% normalized collision energy, and analyzed the fragments in the 200-2000m/z range using 30,000 resolution (m/z = 200). After analysis of the full m/z range (40 DIA scans) a precursor scan was acquired over the 400-1000 m/z range at 60,000 resolution.

#### Peptide library generation

To construct a comprehensive peptide ion library for the analysis of human ovarian cancer we combined several datasets, both internally generated and from external publicly available resources. First, we utilized a publicly available HGSOC proteomics experiment by downloading raw files from the online data repository and searching them through our internal pipeline for data dependent acquisition MS analysis as described in Parker et. al.^21^, and as also performed by others on the same publicly available dataset^22^. To this library, we added an internally generated data set produced by, cation-exchange chromatography (SCX) fractionation of the trypsin digested pooled tumor protein lysates (described above) resulting in 6 fractions. Approximately 1ug of each fraction was loaded onto the Eksigent 415 HPLC system described above and analyzed on a SCIEX TripleTOF 6600 in DDA mode. A 250ms precursor scan was performed on ions between 400-1250 m/z, followed by the isolation and fragmentation of up to the 50 most abundant ions for generation of fragment spectra from 25ms MS2 scans. Spectra generated from these fractions were processed for library generation as described previously^23^Database searches for both the internal and downloaded external datasets utilized human protein sequences defined in a FASTA database of Swiss-Prot-reviewed, Human canonical genome that was downloaded July 2019 and appended with Biognosys indexed retention time (iRT) peptide sequence (Biognosys, Schlieren, Switzerland) and randomized decoy sequences appended. A final, combined consensus spectrast library containing all peptide identifications made between the internal and external dataset was compiled and decoy sequences were appended. The final library has been provided as a supplemental file to this report.

#### Proteomic data analysis

Peptide identification was performed as previously described^23,20^. Briefly, we extracted chromatograms and assigned peak groups using OPENSWATH^24^, utilizing the custom built HGSOC peptide assay library described above and employed PyProphet^25^ to statistically assign a confidence measure (FDR) to each peak group for quality control. TRIC^26^ was used to perform feature-alignment across multiple runs of different samples to reduce peak identification errors. Target peptides with a false discovery rate (FDR) of identification <1% in at least one dataset file, and up to 5% across all dataset files were included in the final results. We used SWATH2stats to convert our data into the correct format for use with downstream software MSstats^27^. The individual patient samples were intensity normalized by dividing the raw fragment intensities by that files total MS2 signal. Dilution series data were not normalized, in order to preserve the intestinal intensity differences induced by the loading schema for that dataset. MSstats was used to convert fragment-level data into peptide- or protein-level intensity estimates via the ‘quantData’ function, utilizing default parameters with the exception of data normalization, which was set to ‘FALSE’.

MS data and supplementary analysis files are deposited on the PASS repository under the identifiers PASS01612 for the dilution series and PASS01613 for the individual patient data set.

### Imputation methods

Missing fragment level intensities were imputed before calculating peptide and protein intensity estimates in MSstats. In the rare cases where a fragment from a given peptide was not detected in any of the samples (N=4 fragments for this dilution dataset), we removed that single fragment from further analysis.

### Zero imputation

This method replaces all missing values with 0.

### minDet

This imputation method replaces all missing values with a minimum value found in the data set. We used the R package imputeLCMD^28^ for this imputation with a q value set to 0.0001.

### minProb

This imputation method performs imputation by random draws from a Gaussian distribution centered in a minimal value which is estimated based on the q^th^ quantile of the observed value. We used the R package imputeLCMD^28^ for this imputation with default parameters.

### Knn (k Nearest Neighbors Imputation)

For missing values, this imputation method finds its nearest values by Euclidean distance and imputes them by averaging the non-missing values of its nearest neighbors. We used impute.wrapper.KNN function from the imputeLCMD^28^ package for this imputation with k = 10 (A range of K values between 10 and 20 have been suggested as appropriate for the knn algorithm ^29,19^).

### SVD (Singular Value Decomposition Imputation)

This method initially replaces all missing values with zero and then estimates them as a linear combination of the k most significant eigen-variables iteratively until it reaches a certain convergence threshold. We used impute.wrapper.SVD function from the imputeLCMD^28^ package in R with the number of PCs set to 2.

### BPCA (Bayesian PCA missing value estimation)

This method combines an expectation-maximization algorithm approach for PCA with a Bayesian model to calculate the likelihood of an estimate for the missing value. We used the R package pcaMethods^30^ with number of PCs used for re-estimation set to 8 i.e., n-1 where n is the number of samples^9^.

### LLS (local least squares imputation)

This imputation method first selects k most similar proteins and then estimates missing values with least squares regression as a linear combination of the values of these k proteins. The similarity is based on the values of samples with complete observations. We used the R package pcaMethods^30^ with parameter k set to 150^31–33^.

### RF (Imputation by Random Forest)

This imputation method uses a robust machine learning algorithm called random forest to build a prediction model. A prediction model is built by defining a target variable with complete observations as the outcome and other variables as predictors. It then iteratively uses the missing values to predict the target variable. The R package missForest^34^ was used with default parameters and parallelize option set to ‘forest’ to improve compute speed.

### Imputation of peptide and protein level missing values

In order to compare the impact of imputation at different levels of data processing in a typical DIA-MS proteomics experiment, we also performed imputation on peptide and protein level data. Imputation at the peptide level was performed by passing the fragment intensities of each sample through the data processing step of MSstats without protein IDs, which forced MSstats to generate an output with intensities for every unique peptide with missing values. These missing values were then imputed with our top three best performing imputation methods, and medians of peptide intensities were used to generate protein intensities for performance evaluation, referred to as peptide level imputation. Non-imputed peptide intensities were used to calculate protein intensities as described above, and the resulting data set was used to impute missing values at the protein level, referred to as protein level imputation.

### Performance evaluation

The performance of each imputation method was evaluated using the metrics described in the recently published R package ‘NAguideR’, as well as by evaluating the impact of imputation methods on the fidelity of our experimentally-induced quantitative differences constructed within the gradient-dataset. Specifically, we calculated the fold changes across each load by taking the difference between mean log2 protein intensities of three replicates across different combinations of loads. Due to the nature of our data set, we expected a log2 fold change of 1 or 2 depending on the loads being compared i.e., fold changes between protein intensities for high load replicates and medium load replicates as well as medium load replicates and low load replicates was expected to be 1; fold changes between protein intensities for high load replicates and low load replicates was expected to be 2. We compared fold changes after imputation to an ideal/expected fold change value of 1 or 2. Pearson’s correlation coefficient was used to calculate the correlation between the fold change of imputed values versus the ideal fold change. The third parameter that we used to evaluate the accuracy of imputation methods was calculating the ratio of Root Mean Square Error (RMSE) to standard deviation of the expected fold changes^11^ (RSR – normalized version of RMSE) compared between imputed fold changes and ideal/expected fold change. R package hydroGOF^35^ was used to calculate RSR values.

Performance evaluation of random forest-based imputation method on the independent data set was done by comparing the set of differentially expressed proteins between primary and recurrent ovarian cancer tumors before and after imputation. ROTS^36^ package was used to perform differential expression analysis with default parameters, including 1000 bootstraps and permutations for resampling. Pearson’s correlation coefficient was also calculated to assess the correlation of replicates before and after imputation.

All evaluations of imputation methods were done using protein intensity values, irrespective of the level at which imputation was performed.

### Other packages used for statistics and figures

All statistical analysis was done in R version 3.6.2^37^. ggplot2^38^, ComplexHeatmap^39^, corrplot^40^, RColorBrewer ^41^ were used to generate figures described in this paper. The schematic for the experimental design of the dilution series experiment was created with BioRender.com.

---

**Supplementary Figure 1.**
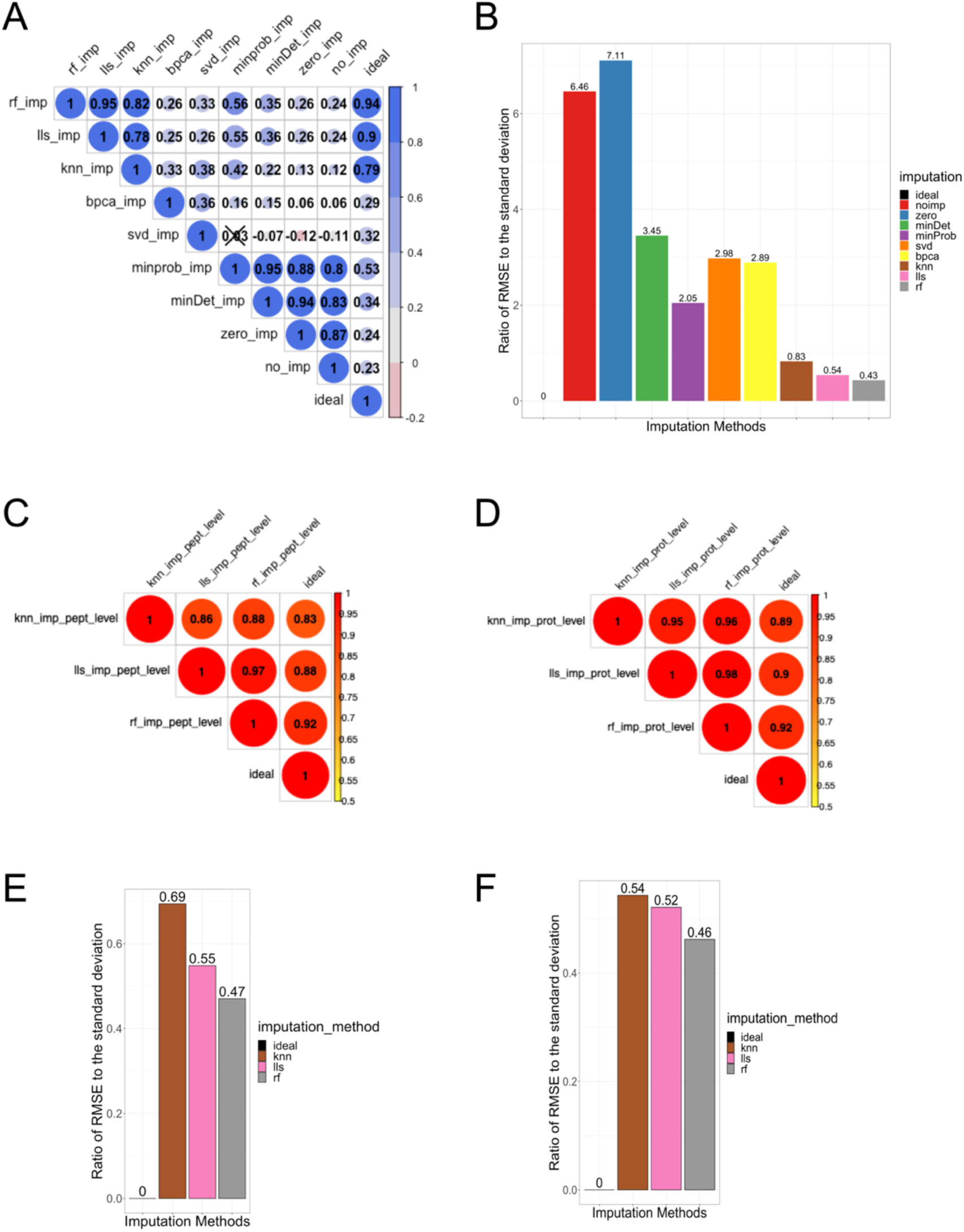
**A)** Pearson’s correlation coefficient shows high correlation between fold change after lls and random forest imputation and ideal/expected fold change (0.9 and 0.94 respectively). **B)** Error estimation between ideal/expected fold change and different imputation methods. **C, D)** Pearson’s correlation coefficient for the top three ranked imputation methods after imputation at the peptide (C) and protein (D) level. **E, F)** Ratio of RMSE to S.D. for the top three ranked imputation methods after imputation at the peptide (C) and protein (D) level

**Supplementary Figure 2:**
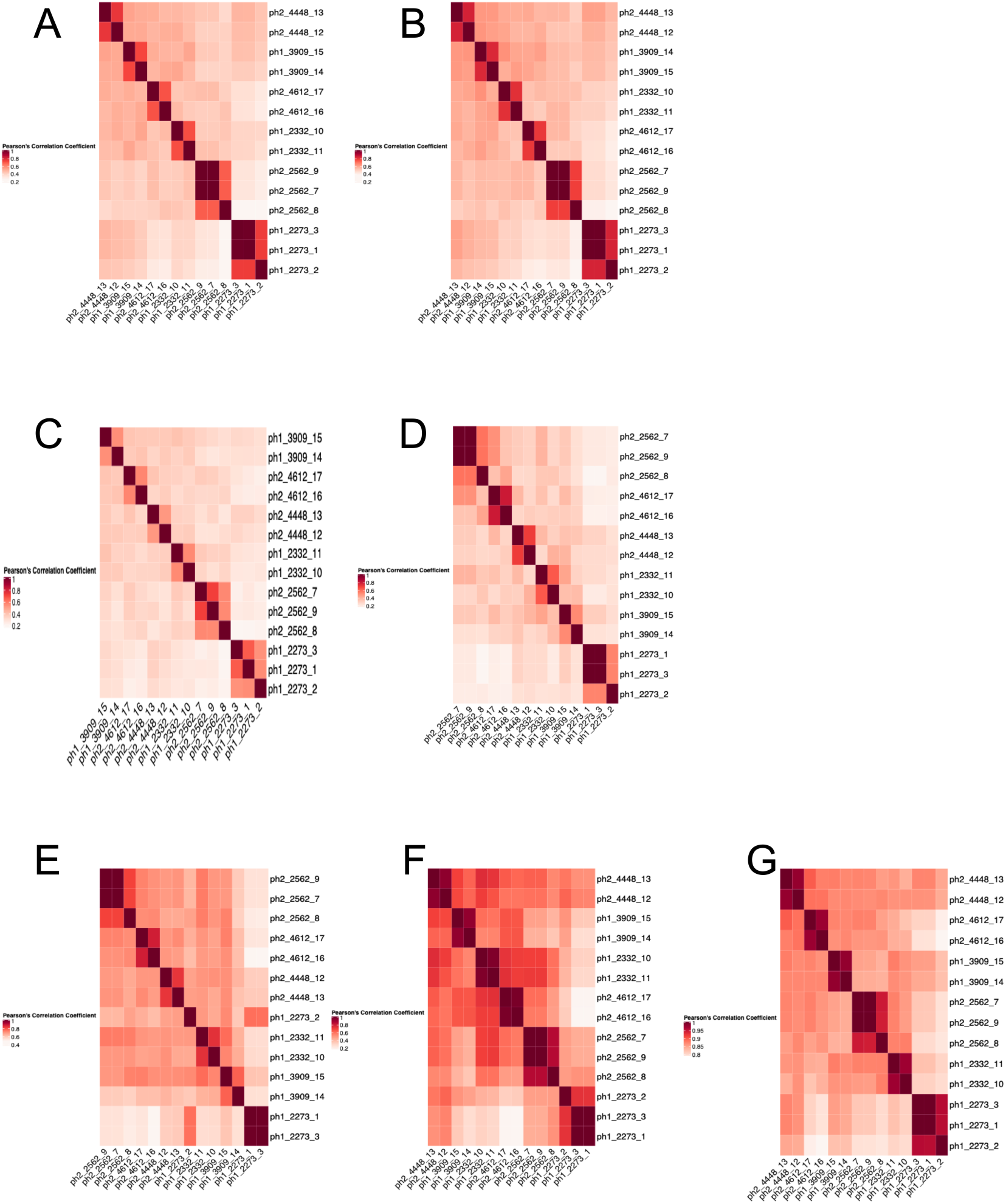
Pearson’s correlation coefficient for replicates of the independent data set by protein intensities before and after imputation of missing values. **A)** Non imputed data set, **B)** minDet imputation, **C)** minProb imputation **D)** SVD imputation **E)** KNN imputation **F)** BPCA imputation **G)** Random Forest imputation

## Notes

### Competing Interest Statement

The authors have declared no competing interest.

